# A gut-microbiota-muscle axis that protects against age-related motor decline by regulating mitochondrial fission in *C. elegans*

**DOI:** 10.1101/2024.12.10.627695

**Authors:** Nathan Dennis, Mireya Vazquez-Prada, Feng Xue, Laura M. Freeman, Antonis Karamalegos, Brigita Kudzminkaite, Ian Brown, Marina Ezcurra

**Author notes:** Correspondence: Marina Ezcurra. These authors contributed equally (Nathan Dennis, Mireya Vazquez-Prada).

## Abstract

Across diverse taxa, the composition of the microbiota is associated with lifelong host health. A mechanistic understanding of how microbial communities influence host physiology could lead to microbiota-based interventions for lifelong health. Here, we have developed a new host-microbiota model system utilising the model organism *C. elegans* combined with a defined natural microbiota (DefNatMta) consisting of 11 bacteria isolated from wild *C. elegans*, to study host-microbiota interactions in a more natural setting. We show that DefNatMta colonises the *C. elegans* gut, forming a stable and distinct gut microbiota. Using DefNatMta, we find a gut microbiota-muscle axis by which the microbiota affects age-related motility and muscular strength and protects against age-related decline in motor function. The gut microbiota-muscle axis acts by altering metabolism and mitochondrial network dynamics in muscle, and requires dynamin-related protein 1 DRP-1, a regulator of mitochondrial fission to protect against age-related motility decline. Our study demonstrates a gut microbiota-muscle axis and microbiota-mitochondria communication affecting age-related muscle function.

## Introduction

The gut microbiota forms a complex ecosystem impacting host metabolism and homeostasis throughout life. Across diverse taxa, the microbiota is associated with host development ^1^, metabolism ^2^ and health ^3^. The gut microbiota is a key factor in multiple functions such as energy/nutrient absorption, immunity, intestinal permeability, hormone, and vitamin production and microbial functions influence health and fitness of their hosts. Gut microbes produce a vast number of enzymes and bioactive products and that can influence health; some are beneficial, but others have toxic effects. Consequentially, alterations of the composition of the microbiota can lead to signalling to host tissues, contributing to altered host metabolism, dysregulated bodily functions, diseases, ageing and frailty ^4–7^.

The molecular mechanisms by which host-microbiota interactions affect the function of different tissues across life stages are not well understood. This is at least in part due to the high level of complexity of the human microbiota, consisting of hundreds of species altered by diet, lifestyle and genetics. Moreover, species in the human microbiota are often anaerobic, difficult to cultivate and require growth conditions only found in the gut ^8^, further limiting a mechanistic understanding. Laboratory studies using mice have improved the understanding of causative relationships between the microbiota and host metabolism and health, but often involve using germ-free mice, requiring costly and labour-intensive specialised facilities and equipment, and only testing a limited number of health outcomes at single timepoint ^3^. A better understanding of how microbial communities influence host health could lead to microbiota-based interventions to promote lifelong health.

*C. elegans* is a simple tractable model organism that offers informative and cost-effective routes to establish mechanistic understanding of host-microbiota interactions. Cellular functions and metabolic pathways fundamental for health and ageing are evolutionary conserved between *C. elegans* and mammals, making *C. elegans* is ideal to understand how microbiotas impact on host health across a range of health measures and life stages. To perform mechanistic studies of host-microbiota interactions affecting health and fitness, we have established a host-microbiota model using *C. elegans* combined with a simple microbial community. The community consists of 11 bacterial strains isolated from wild *C. elegans* (DefNatMta), reflecting the taxonomic diversity of their native microbiota ^9–11^. The strains are aerobic and grow in standard laboratory conditions.

Here, we establish DefNatMta combined with *C. elegans* as a highly tractable system to study natural host-microbiota interactions and determine the underlying mechanisms. We find that DefNatMta stably colonises the *C. elegans* intestinal tract, and specific microbiota species are enriched within the gut. DefNatMta sustains *C. elegans* growth, development and reproduction and is suitable for laboratory cultivation. We show that DefNatMta affects age-related motility and strength in the host, reducing motility in young adults but protecting against age-related decline, demonstrating a microbiota-muscle axis affecting age-related motor function. We further demonstrate microbe-mitochondrial communication by which DefNatMta alters mitochondrial networks to protect against age-related decline in motility.

## Results

### DefNatMta colonises *C. elegans* and forms a stable and distinct gut microbiota

To characterise host-microbiota interactions in *C. elegans* we established a defined natural microbiota (DefNatMta) consisting of 11 bacterial isolates identified in wild *C. elegans* (Fig. 1A). The bacterial isolates were selected based on previous work examining the gut contents of wild *C. elegans* samples isolated from compost, rotting fruit and invertebrate vectors ^9^. This defined microbiota contains some of the most abundant bacterial species found within these wild *C. elegans* samples, which together constitute a taxonomically diverse community reflective of the native *C. elegans* microbiota. To study the impact of this community on the worms, all bacterial isolates were cultivated in LB at 25°C, mixed in equal volumes, and seeded onto NGM plates, using *E. coli* OP50 as control (Fig. 1B). OP50 is widely used as the standard laboratory diet for *C. elegans* cultivation but is a poor coloniser of young healthy *C. elegans*, due to grinding of the bacteria by the pharynx, and host immunity, reducing bacterial load and proliferation in the intestine (Portal-Celhay, Bradley and Blaser, 2012; Cabreiro and Gems, 2013). We verified whether DefNatMta colonises the *C. elegans* gut by examining the number of viable cells recoverable from the gut contents of worms. Colony forming units (CFU) from the gut contents of DefNatMta-fed worms were 12-fold higher than those of OP50-fed worms on day 1 of adulthood (Fig. 1C). The difference in CFU count was maintained up to day 7 of adulthood, showing that in contrast to OP50, DefNatMta forms a stable bacterial community in the *C. elegans* gut. We additionally assessed colonisation dynamics using 16s rRNA sequencing, comparing the gut contents of DefNatMta-fed worms to the composition of the DefNatMta lawn. Our results demonstrated the formation of a selective gut microbiota, compositionally distinct from the bacterial lawn (Fig. 1D). Principal coordinates analysis of Bray-Curtis dissimilarities confirmed this distinction (Fig. 1E; MANOVA PCo1-PCo3: P *<* 0.01), indicating that the major taxa involved were *Ochrobactrum* MYb71 (highly enriched in the gut), *Acinetobacter* MYb10 (depleted in the gut relative to the lawn), and *Stenotrophomonas* MYb57 (moderately enriched in the gut), consistent with other studies showing intestinal colonisation by MYb71 and MYb57 ^9,14^. Despite the difference in microbial composition between the gut and the lawn there was no change in α-diversity between gut content and bacterial lawns (Fig. S1).

**Figure 1:**
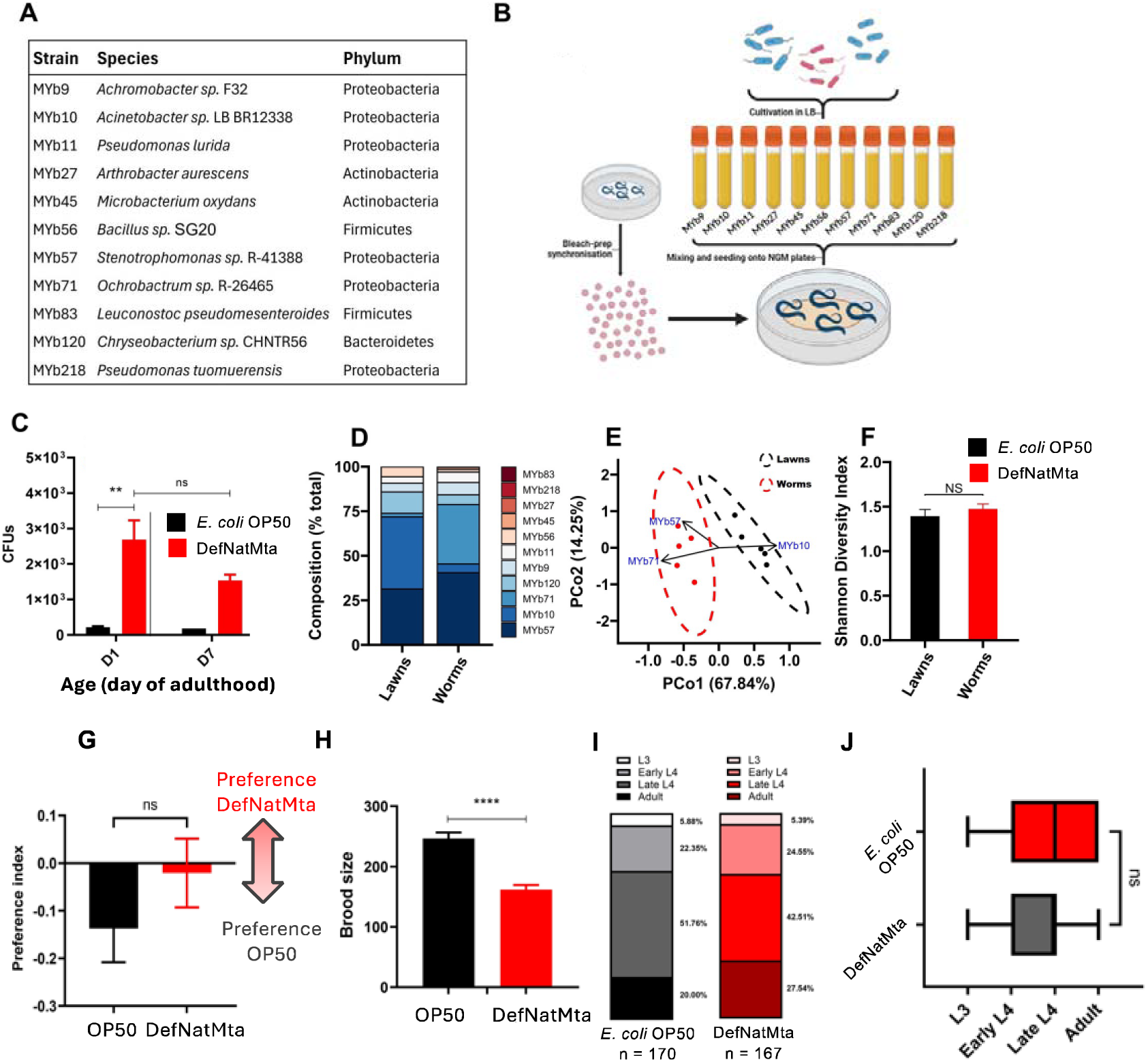
DefNatMta colonises C. elegans and sustains growth. **A**) Composition of DefNatMta with phylogenetic details. Bacteria were sourced from wild C. elegans samples isolated from compost, rotting fruit or invertebrate vectors by the Schulenberg lab Dirksen et al. (2016). **B**) Schematic of experimental set-up. **C**) CFUs recovered from the gut contents of animals fed with the experimental microbiota mixture or E. coli OP50. n = 150 per condition, pooled from 3 biological experiments. **D**) Species-level composition of the experimental microbiota lawn and the gut contents of experimental microbiota-fed worms. N = 5. **E**) Principal coordinates analysis of microbial diversity from experimental microbiota lawns and experimental microbiota-fed worms; PCoA was performed using Bray-Curtis dissimilarities. The ellipses represent the 95% CIs, and the arrows represent the taxa significantly correlated with the ordination axes (P < 0.05; lengths scaled by R2). Arrows were fitted using the envfit function for vegan v. 2.6-6.1. **F**) Shannon diversity indices of experimental microbiota lawns and experimental microbiota-fed worms. **G**) Food choice behaviour of day 1 adult worms reared on either E. coli OP50 or the experimental microbiota. Indices > 0 correspond to preference for the experimental microbiota; indices < 0 correspond to preference for E. coli OP50. N = 11-12 repeats, pooled from 3 experiments. Total animals tested was 705 for DefNatMta and 1164 for OP50. **H**) Total brood size of experimental microbiota and E. coli OP50-fed worms. n = 22-29 pooled from 3 experiments. **I-J)** Distributions of developmental stages (L3-Adult) 60h after egg preparation synchronisation. n = 167-170 pooled from 3 experiments. Statistical analysis performed using a Wilcoxon rank-sum test. Not significant (P > 0.05). Unless otherwise stated, data were analysed using t-tests. ****, P < 0.0001; **, P < 0.01*; P < 0.05; ns, not significant (P > 0.05).

After establishing the formation of a unique gut microbiota, we confirmed that DefNatMta is suitable for cultivation of C. elegans in the laboratory. DefNatMta-fed animals produced an average of 162 progeny by self-fertilisation, a reduction compared to 247 for animals fed with OP50. Developmental timing to reach adulthood was not affected, indicating that the DefNatMta provides adequate nutrition to developing worms (Fig. 1I, J). In a food choice assay, animals did not show preference for neither OP50 nor DefNatMta, showing that DefNatMta is not perceived as an inferior or pathogenic food source (Fig. 1F). These results demonstrate that DefNatMta forms a stable, abundant and selective community within the gut suitable for cultivation of *C. elegans* in the laboratory.

### The DefNatMta protects against age-related decline in muscle function

To further test how DefNatMta affects host health, we measured movement using a swimming assay, a key indicator of locomotive behaviours and body wall muscle function, combined with automated image analysis (Fig 2A). DefNatMta reduced swimming rate in young adults by 14.6% but protected against age-related decline in muscle function. Like ageing humans, *C. elegans* undergo sarcopenia, a progressive decline in muscle mass and strength with age, accompanied by reduced muscle function and motility ^15,16^. Consistent with previous studies, we found that in animals fed with OP50, swimming rate declined by 30% between days 1 and 7 of adulthood, while animals fed with DefNatMta did not undergo any significant decline during this time interval (age x diet interaction: P*<*0.0001). DefNatMta-fed animals instead exhibited an increase in thrashing rate at day 4 of adulthood, retaining 90% of their day 1-adult thrashing rate at day 7 of adulthood (Fig. 2B, C). Stretch, a measure of the average depth of thrashing ^17^, was significantly increased by DefNatMta on days 1 and 7 of adulthood, indicative of improved motor function (Fig. 2D). Further assessment of muscle strength in three-dimensional media using Pluronic gel burrowing assays ^18^ generated results similar to the thrashing assays with DefNatMta reducing burrowing in young adults but protecting against age-related decline (Fig. 2E). We asked if DefNatMta also affects other measures of motor function and measured execution of the defecation motor program, a series of coordinated muscle contractions that occur in a cycle to expel intestinal contents, and pharyngeal pumping, a rhythmic, neuromuscular process that allows *C. elegans* to feed. DefNatMta did not alter defecation rates but increased pharyngeal pumping in day 1 adults (Fig 2F, G). These findings suggest a microbiota-muscle axis acting on body wall muscle to affect muscle functionality and protect against age-related decline in motility.

**Figure 2:**
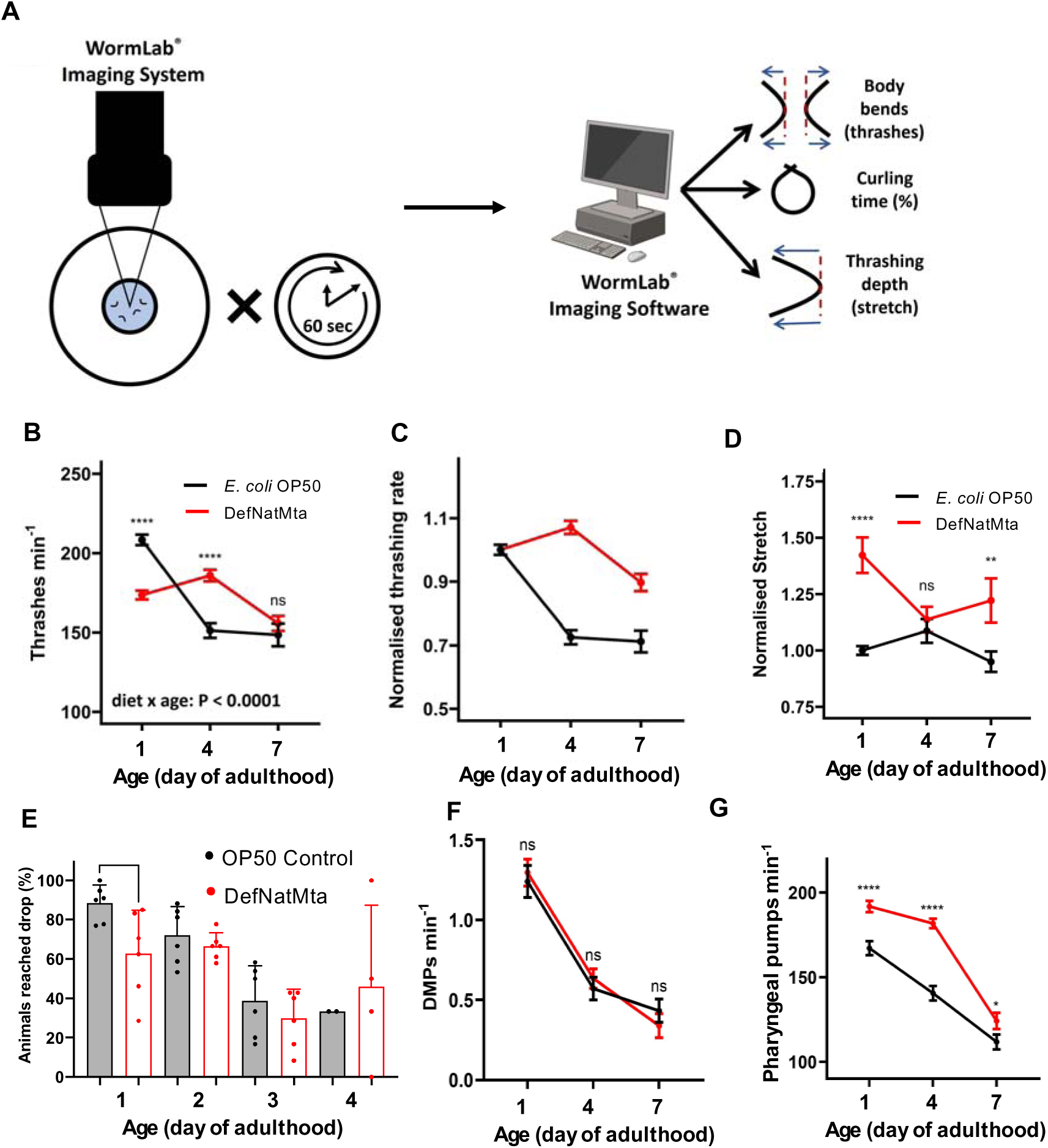
DefNatMta alters host motility and muscle function. **A**) Schematic of motility assessment using the WormLab^©^ Imaging System and Software. **B-C**) Raw (B) and day 1 adult-normalised (C) frequencies of lateral swimming (thrashing rate) of OP50 and DefNatMta-fed worms. n = 24-31 pooled from 2 experiments. **D**) Thrashing depth (stretch; normalised to mean values of D1 OP50-fed worms. n = 24-31 pooled from 2 experiments. ****, P *<* 0.0001; **, P *<* 0.01; *, P *<* 0.05; ns, not significant (P *>* 0.05). **E**) Burrowing performance of OP50 and DefNatMta-fed worms at 120 mins in PF-127 gel. n = 78 per condition pooled from 2 experiments with three technical repeats each. **F**) Defecation rate (DMP: defecation motor programme) of experimental microbiota and E. coli OP50-fed worms. n = 30 pooled from 3 experiments. **G**) Pharyngeal pumping rate of experimental microbiota and E. coli OP50-fed worms. N = 90-120 pooled from 3 experiments.

### DefNatMta alters host mitochondrial dynamics in body wall muscle

To understand the mechanisms by which gut-microbiota-muscle axis affects muscle function we first tested if DefNatMta protects against the breakdown of sarcomere organisation ^16,19^ or neuromuscular junction function that occurs with age ^20^. Assessment of the body wall musculature using electron microscopy and quantification of sarcomere organisation using fluorescently tagged myosin and actin in body wall muscle ^21^ did not show any differences between OP50- and DefNatMta-fed animals (Fig. S2). Also examination of neuromuscular transmission efficiency showed no microbiota-induced effects. There were no differences between DefNatMta- and OP50-fed worms in contractability or response to touch stimuli ^22,23^ (Fig. S2).

The lack of association between DefNatMta and host body wall musculature and neuromuscular transmission led us to examine if DefNatMta mediates effects on age-related muscle function by altering mitochondrial function. Age-related alterations of mitochondrial morphology have been described across species^24–26^ and *C. elegans* exhibit well-described age-related alterations in mitochondrial morphology and networks in muscle ^27^. Using GFP-tagged mitochondria in body wall muscle^28^, we found that the structure of mitochondrial networks became more fragmented with age in both OP50- and DefNatMta-fed animals (Fig. 3A-C). DefNatMta significantly increased mitochondrial fragmentation on days 4, 7 and 11 of adulthood compared to OP50 (Fig. 3B, C), suggesting host-microbiota interactions that alter mitochondrial networks in muscle.

**Figure 3:**
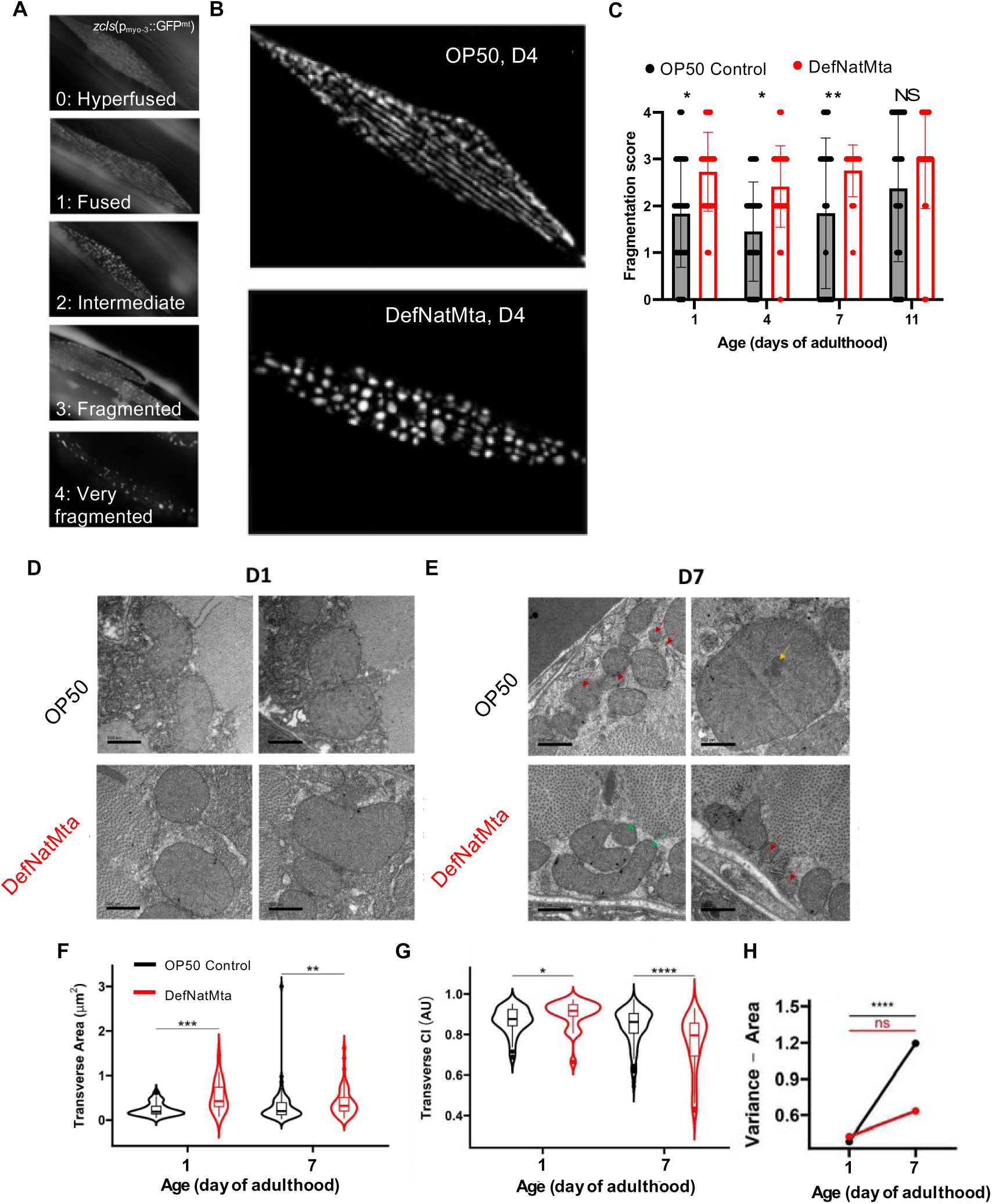
DefNatMta alters mitochondrial morphology and ultrastructure in body wall muscle. **A**) Representative images of discrete numerical scores describing mitochondrial fragmentation in *zcIs14(myo-3::GFP^mt^)* animals. 0: Mitochondria are hyperfused; tubular networks of mitochondria are joined; 1= Linear: tubular networks of mitochondria organised into linear tracts; 2 = Intermediate: both tubular and spherical mitochondria present in cell; 3 = Fragmented: mostly spherical mitochondria covering majority of cell; 4 = Very fragmented: large regions of cell devoid of GFP^mt^ signal. **B**) Representative images of mitochondrial morphology in body-wall muscle in *zcIs14(myo-3::GFP^mt^)* animals. Tubular networks in OP50-fed (top) and fragmented networks in DefNatMta-fed (bottom) animals at day 4 of adulthood. **C**) DefNatMta-fed animals exhibit higher mitochondrial fragmentation scores on days 1, 4 and 7 of adulthood. Data points show fragmentation score for each animal. Bars show mean fragmentation scores. N ≥ 36 pooled from 3 experiments. Statistical analysis by Chi-square test. **D-E**) Representative micrographs of mitochondria from transverse sections of the body-wall muscle of OP50- and experimental DefNatMta-fed worms during day 1 (E) and day 7 (F) of adulthood. Mitochondria with clear outer membrane dysfunction (red arrows), electron-dense inclusions (yellow arrows) and unusually low circularity indices (green arrows) are highlighted. Scale bars, 500 nm. **F-G**) Mitochondrial area (G) and mitochondrial circularity (CI; H) calculated from transverse sections represented in D-E. Mitochondrial ultrastructure and morphology was measured in all observable mitochondria per transverse section: n = 23-129 pooled from 2 experiments. Significance calculated using Wilcoxon rank-sum tests. **H**) Mitochondrial area variance calculated from transverse sections represented in Fig. 3E-F. Mitochondrial ultrastructure and morphology were measured in all observable mitochondria per transverse section: n = 23-129 pooled from 2 experiments. Significances presented represent variance comparisons between day 1 and day 7 using Fisher’s F-tests. ****, P < 0.0001; ***, P < 0.001; **, P < 0.01; ns, not significant (P > 0.05).

Examination of mitochondrial morphology and ultrastructure using electron microscopy showed a uniform internal mitochondrial structure with fully intact and regular outer membrane morphologies and dense cristae in both conditions on day 1 of adulthood (Fig 3D). In day 7 adults, mitochondrial abnormalities were visible, including lack of a discernible outer membrane around all or part of the mitochondrion, sparse cristae, abnormally large mitochondria, and mitochondria with electron dense inclusions (Fig. 3E). Mitochondria of day 1 adults fed on DefNatMta were larger and more circular than those of OP50-fed worms, consistent with a globular, fragmented mitochondrial network (Fig. 3D, F, G). In day 7 adults the difference in the size of the mitochondria remained, but the mitochondria of DefNatMta-fed worms had a significantly lower average circularity index than those of OP50-fed worms, (Fig. 3E-G). Additionally, the variation in mitochondrial morphology was altered by DefNatMta. The variance in mitochondrial area increased 3.2-fold in OP50-fed worms between day 1 and day 7 of adulthood but did not significantly increase in DefNatMta-fed worms, while the variance of circularity indices significantly increased in both OP50-fed worms and DefNatMta-fed worms (Fig. 3H), suggesting the DefNatMta reduces variability in mitochondrial area but not circularity. Together these results provide evidence that the DefNatMta influences host mitochondrial networks, altering mitochondrial size and circularity, without affecting mitochondrial ultrastructure.

### Microbiota-induced motility preservation requires regulation of mitochondrial fission dynamics

Given that DefNatMta both alters mitochondrial networks and motility, we explored if these effects are linked. We first examined if there was a correlation between mitochondrial fragmentation and motility. Animals with GFP-tagged muscle mitochondria were aged and classified according to their ability to move (A: highly mobile, spontaneous sinusoidal locomotion; B: mobile only when prodded; C: alive but immobile even when prodded)^16^. Each animal was then imaged individually, and mitochondrial fragmentation severity was scored. This allowed us to determine the motility class and mitochondrial fragmentation score of each individual. As expected, mitochondrial fragmentation scores were positively correlated with impaired motility in OP50-fed animals (Fig 4A). In contrast, no association between motility class and mitochondrial fragmentation score was found in DefNatMta-fed worms, suggesting DefNatMta uncouples mitochondrial network organisation from age-related muscle function.

**Figure 4:**
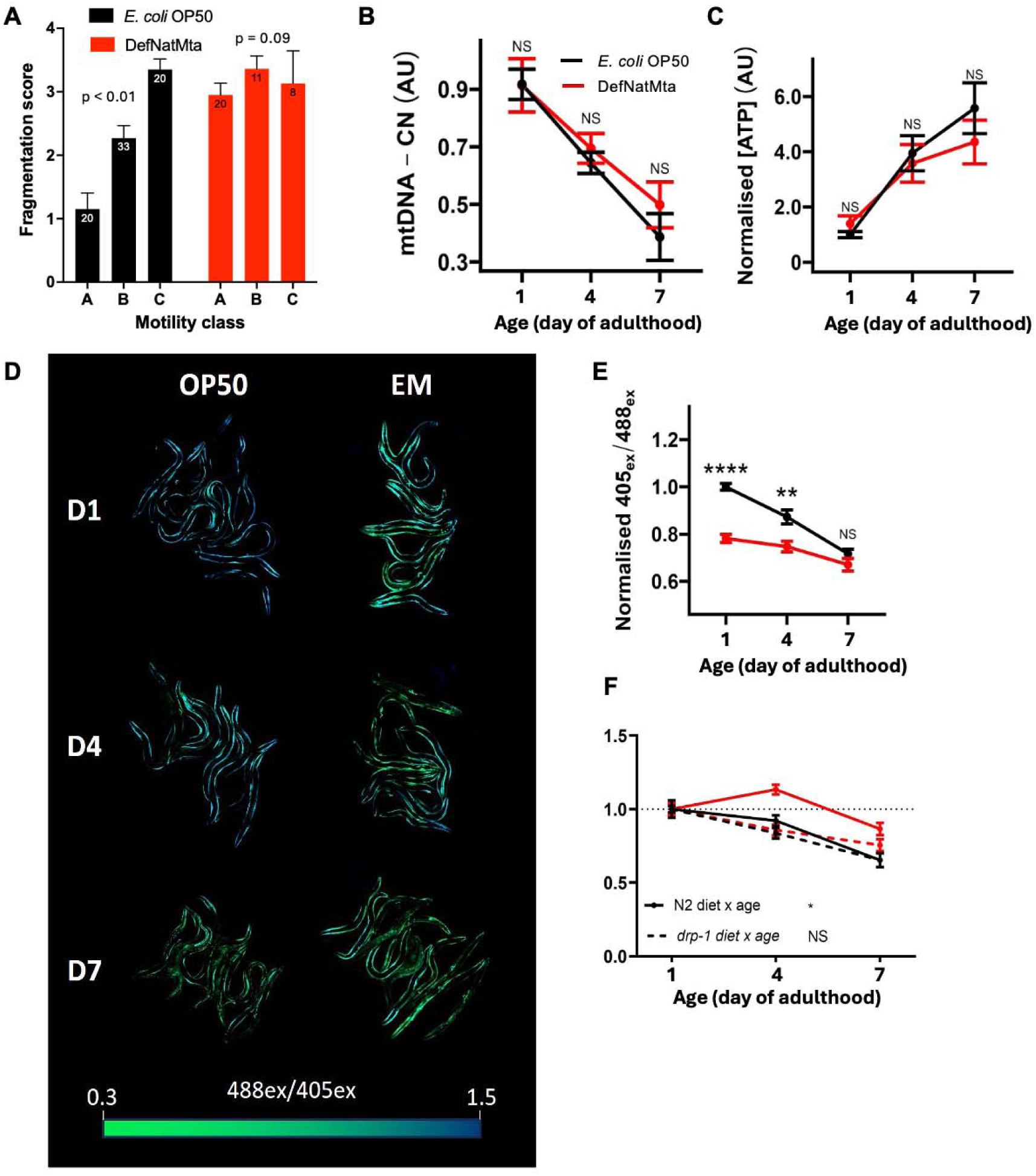
DefNatMta alters muscle ATP levels and requires mitochondrial fission to preserve age-related motility. **A**) Increased mitochondrial fragmentation scores are correlated with reduced motility in aged OP50-fed animals. The correlation is lost in DefNatMta-fed animals. Motility classes: A = Highly mobile, spontaneous movement; B = Mobile when prodded but no spontaneous movement; C = Alive but not mobile. Statistical analysis by Chi-square test assessing independence of fragmentation score and motility class. Bars represent mean fragmentation scores per motility class with numbers of animals indicated within. n = 39-73 pooled from 3 experiments. **B**) Whole-body mitochondrial DNA copy-number (mtDNA-CN) measured in single animals in *glp-1* background by qPCR. Relative mtDNA-CNs were calculated comparing amplification of mitochondrial and nuclear DNA and normalised to average relative mtDNA-CNs of relevant OP50-fed controls. n = 11-13 pooled from 2 experiments. **C**) Whole-body ATP levels quantified using a luminescence assay and a standard curve of ATP standards and normalised to average ATP levels of relevant OP50-fed controls.. n = 15-23 pooled from 4 experiments. **D**) Representative full body scans of animals expressing a Queen-2M ratiometric ATP sensor in the body wall muscles. Images are saturated for visualisation. **E**) Normalised 405*_ex_* / 488*_ex_* ratios calculated from raw fluorescence intensities of whole animals as represented (C). N = 35-43 pooled from 3 experiments. **F**) Normalised thrashing rates (relative to day 1 adults) of OP50- and DefNatMta fed worms with loss of function mutation in the mitochondrial fission gene *drp-1* (*tm1108*; G). n = 16-56 pooled from 3 experiments. Unless otherwise stated statistical analysis performed using analysed by ANOVA with age and diet included as an interaction term; ****, P *<* 0.0001; **, P *<* 0.01; *, P *<* 0.05; ns, not significant (P *>* 0.05).

We asked if the effects DefNatMta on mitochondrial networks was reflected in readouts of mitochondrial function and measured whole-body mtDNA and ATP levels as indicators of mitochondrial health^29^. Both ATP and mtDNA levels were unaffected by the DefNatMta (Fig. 4B, C), suggesting that DefNatMta-induced changes to mitochondrial network morphologies are not reflected in altered ATP or mtDNA content at the whole-body level. Next we examined ATP levels in body wall muscle specifically using the ratiometric ATP sensor Queen-2M under the control of the *myo-3* promoter^30,31^. The effects of DefNatMta on muscle ATP levels mirrored the effects we observed on swimming; DefNatMta-fed worms had significantly lower ATP levels relative to OP50-fed worms in early adulthood but exhibited less age-related decline (age x diet interaction: P*<*0.01), and had similar ATP levels to OP50-fed worms by day 7 (Fig. 4D-E). These findings show that DefNatMta alters muscle bioenergetics specifically.

To better understand the mechanisms by which DefNatMta affects muscle function, we directly tested if mitochondrial network dynamics are involved in the motility phenotype utilising mutants for *drp-1*, which encodes Dynamin-related protein 1. *drp-1* is required for mitochondrial fission and *drp-1* mutants exhibit hyperfused mitochondrial networks ^32^. Thrashing assays in *drp-1* mutants showed similar rates of age-related motility decline regardless of diet (Fig. 4F). Our findings demonstrate microbe-mitochondria communication by which DefNatMta alters muscle bioenergetics and mitochondrial networks, specifically implicating mitochondrial fission in the motility-preserving effects of DefNatMta.

## Discussion

The mechanisms by which the microbiota impacts on host lifelong health remain largely unknown. Using the genetically tractable model organism *C. elegans* combined with a microbial community derived from the natural *C. elegans* microbiota, DefNatMta, we found that this community affects age-related motility in the host. Using this model system, we identified underlying mechanisms and demonstrated a microbiota-muscle axis by which DefNatMta impacts on muscle ATP levels, mitochondrial networks, and motor function. We showed that DefNatMta requires mitochondrial fission to protect against age-related decline in motility, directly demonstrating microbe-mitochondrial communication involving regulation of mitochondrial fission underlying the microbiota-muscle axis.

Our study highlights mechanisms by which the gut microbiota influences muscle health during ageing. In humans individual differences in gut microbiota are linked to muscle function and frailty and changes in microbiota composition are correlated with frailty and age-related muscle loss ^33,34^ Studies using germ-free and antibiotic treated mice have provided experimental evidence that the gut microbiota regulates skeletal muscle mass and function ^35,36^. Our study supports and add to these findings by showing that the microbiota alters mitochondrial networks and metabolism in muscle, thereby providing mechanistic insight into the gut microbiota-muscle axis. We anticipate that future studies will add more mechanistic detail, for example the gut microbiota is likely to alter key biological processes such inflammation, nutrient bioavailability, and lipid metabolism, that contribute to mitochondrial metabolism and muscle function ^37^.

The gut microbiota is complex, making experimental manipulation and demonstrating causation challenging. Using a defined microbiota consisting of bacterial species isolated from wild *C. elegans* allows us to perform mechanistic studies in a more natural setting. We find that in comparison to *E. coli* OP50, a domesticated bacteria and artificial *C. elegans* diet, DefNatMta forms a stable gut microbial community. The formation of this gut microbiota is characterised by the enrichment of *Ochrobactrum vermis* MYb71 and *Stenotrophomonas* MYb57, consistent with previous studies ^9,14^. DefNatMta contains many genera featured in a recently developed *C. elegans* microbiota resource, CeMbio^38^. Similarly to CeMbio, DefNatMta enables more realistic studies whereby natural host-microbiota interactions are taken into account.

## Acknowledgements

We acknowledge support from the Biotechnology and Biological Sciences Research Council (BBSRC) through grant BBSRC(BB/V011243/1) and a SoCoBio studentship awarded to Laura Freeman, and Dr John Stolz through a PhD studentship awarded to Nathan Dennis. Some strains were provided by the CGC, which is funded by NIH Office of Research Infrastructure Programs (P40 OD010440). We thank Jennifer Tullet and Nazif Alic for meaningful discussions on the manuscript.

## Materials and Methods

### Bacterial culture and strains

Strains *Achromobacter sp.* F32 MYb9, *Acinetobacter sp.* LB BR12338 MYb10, *Pseudomonas lurida* MYb11, *Arthrobacter aurescens* MYb27, *Microbacterium oxydans* MYb45, *Bacillus sp.* SG20 MYb56, *Stenotrophomonas sp.* R-41388 MYb57, *Ochrobactrum sp.* R-26465 MYb71, *Leuconostoc pseudomesenteroides* MYb83, *Chryseobacterium sp.* CHNTR56 MYb120 and *Pseudomonas tuomuerensis* MYb218^9^ were used to assemble DefNatMta. The strains were grown individually in a 25°C for three days before being mixed in equal volumes to form the DefNatMta mixture. 200 µL aliquots were seeded onto 60 mm NGM plates and allowed to dry at room temperature for three days. *E. coli* OP50 was grown at 37°C in a 180 RPM shaking overnight prior to seeding plates.

### *C. elegans* culture and strains

*C. elegans* were maintained under standard conditions^39^, at 20°C on NGM plates seeded with *Escherichia coli* OP50. All experiments were conducted at 20°C unless otherwise stated. For sterilisation and synchronisation, *C. elegans* were treated with sodium hypochlorite/sodium hydroxide, followed by aliquoting eggs onto experimental plates.

The following strains were used: N2 (wildtype N2 male stock, N2 CGCM) [55] GA2001 *wuls305 [myo-3p::Queen-2m]*, CB4037 *glp-1 (e2141)*, RW1596 *myo-3 (st386); StEx30 [myo-3p::GFP::myo-3 + rol-6 (su1006)*, WX8490 *YqIs100 [ced-1p::mCHERRY::ACT1]*, SJ4103 *zcls14 [myo-3p::GFP(mit)* and CU6372 *drp-1 (tm1108)*.

### Gut colonisation

Gut colonisation was assessed by counting the number of colony-forming units (CFUs) recoverable from nematode samples following surface sterilisation. For each assay, 50 animals were anaesthetised in 25 mM tetramisole in M9, surface-sterilised with a bleach solution (2.5% sodium hypochlorite, 2M NaOH) for 5 minutes, washed three times with M9, and homogenised in 1 % Triton X-100 using a pellet pestle. The resulting suspensions were then centrifuged at 13 000 x g for 10 minutes, resuspended in M9, diluted at 10^-1^, 10^-2^ and 10^-3^ in sterile water, and spread onto LB plates in technical triplicate. CFUs were then counted following either 24 h incubation at 37°C (*E. coli* OP50) or 72 h incubation at 25°C (DefNatMta), and normalised to the number of worms as follows:

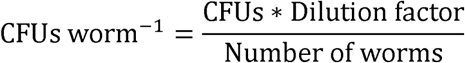

### DNA extraction for microbiome composition analysis

Genomic DNA from *C. elegans* guts was extracted from worm lysates based on a modified version of a previous protocol^14^. Briefly, populations of worms were collected into Eppendorf tubes, pelleted by centrifugation at 2500 RPM for 30 seconds, and washed twice with M9. The animals were surface-sterilised (2.5 % sodium hypochlorite, 2M NaOH) for 5 minutes, and washed a further three times with M9. The efficacy of surface sterilisation was checked by examining CFU counts recoverable from the supernatant immediately following the final wash (500 µL aliquots were spread on LB plates and incubated for 24 h at 25 °C), and only samples showing CFU counts of zero were sequenced. DNA was extracted using a QIAGEN DNeasy Blood and Tissue Kit and subject to three freeze-thaw cycles at −80 °C and 37 °C to release intestinal bacterial cells into solution. DNA purity and yield were assessed using a NanoDrop UV/Vis spectrophotometer and a Qubit Fluorometer, respectively.

Genomic DNA from bacterial lawns were extracted in tandem to gut extractions. Contents of three three-day old lawns seeded on NGM plates were scraped into 2 mL M9 using an inoculating loop. DNA extraction and quantification followed the same protocol outlined above.

### 16s rRNA sequencing

Sequencing of genomic DNA was performed by Novogene UK Ltd. Bacterial diversity was examined by PCR amplification of the V4 region of 16s rRNA with barcoded primers (515F-806R)^40^. PCR amplicons were quantified with SYBR Green, pooled in equidensity ratios, and purified using a QIAGEN Gel Extraction Kit. Library preparation was performed using an NEBNext Ultra DNA Library Prep Kit, and sequencing with an Illumina paired-end platform with a read length of 250 bp. Paired-end reads were merged with FLASH v. 1.2.7^41^. Quality filtering was conducted using QIIME v. 1.7.0^40^ with default settings. Chimeric sequences were removed using the UCHIME algorithm with the Gold reference database^42^. Operational taxonomic units (OTUs) were constructed using Uparse v. 7.0.1001^43^ with a minimum percent identity of 97 %. Species assignment of representative OTU sequences was conducted using the Ribosomal Database Project Classifier v. 2.2^44^. OTU sequences failing to match published DefNatMta sequences were removed (< 5 % of all reads). OTU abundances were normalised using a standard sequence number corresponding to the sample with the fewest sequences.

### Microbiome composition analysis

All diversity analyses were performed with the *vegan* package for R v. 4.3.3 using normalised OTU abundances. Alpha diversity was assessed using a Shannon diversity index. Beta diversity was assessed via principal coordinates analysis (PCoA) of Bray-Curtis distances using the *capscale* function. The relationships between environmental variables and bacterial diversity were assessed using the function “*envfit*” with 999 permutations. *envfit* computes the direction of the effects of continuous variables (here taxon abundances) by calculating the direction of maximum correlation between a given continuous variable and the ordination scores, and the average ordination score for each level of each categorical variable (here sample source). The statistical significances of these variables are then assessed using a permutation test.

### Food preference

Food-choice behaviour was assessed using a modified version of a previous protocol^45^. Worms were collected in groups of at least 20, washed three times with M9, and transferred onto the centres of 30 mm NGM plates seeded with 15 µL lawns of *E. coli* OP50 and the DefNatMta mixture, with each lawn spaced 20 mm apart. The plates were then incubated at 20°C for 24 h and imaged with an MBF Biosciences WormLab Imaging System. The number of worms residing in each bacterial lawn were counted by eye and a food preference index was calculated as follows:

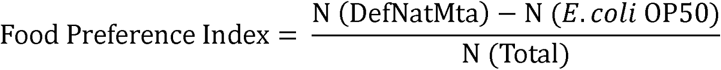

With this index, a value of 1 occurs when all animals associate with the DefNatMta mixture, and a value of −1 occurs when all animals associate with *E. coli* OP50.

### Brood size

Brood size was assessed as described^46^. Briefly, individual worms were transferred at the L4 stage to separate NGM plates. Subsequently, they were transferred each day, up to day 6 of adulthood, to freshly seeded NGM plates to lay eggs. The viable progeny hatching from each brood were then counted at the late larval to adult life stage.

### Developmental rates

Developmental rates were assessed via light microscopy 60 h after age synchronisation (see *Bacterial culture* and strains

Strains *Achromobacter sp.* F32 MYb9, *Acinetobacter sp.* LB BR12338 MYb10, *Pseudomonas lurida* MYb11, *Arthrobacter aurescens* MYb27, *Microbacterium oxydans* MYb45, *Bacillus sp.* SG20 MYb56, *Stenotrophomonas sp.* R-41388 MYb57, *Ochrobactrum sp.* R-26465 MYb71, *Leuconostoc pseudomesenteroides* MYb83, *Chryseobacterium sp.* CHNTR56 MYb120 and *Pseudomonas tuomuerensis* MYb218^9^ were used to assemble DefNatMta. The strains were grown individually in a 25°C for three days before being mixed in equal volumes to form the DefNatMta mixture. 200 µL aliquots were seeded onto 60 mm NGM plates and allowed to dry at room temperature for three days. *E. coli* OP50 was grown at 37°C in a 180 RPM shaking overnight prior to seeding plates.

### *C. elegans* culture and strains)

Nematodes were anaesthetised in 25 mM tetramisole and mounted onto 2.5 % agarose pads in groups of approximately 50. Each animal was observed using a Leica DMR compound epifluorescence microscope with a 40X objective and scored based on vulval structure and the presence or absence of oocytes as described ^47^, see Table 1 for scoring criteria.

**Table 1.**
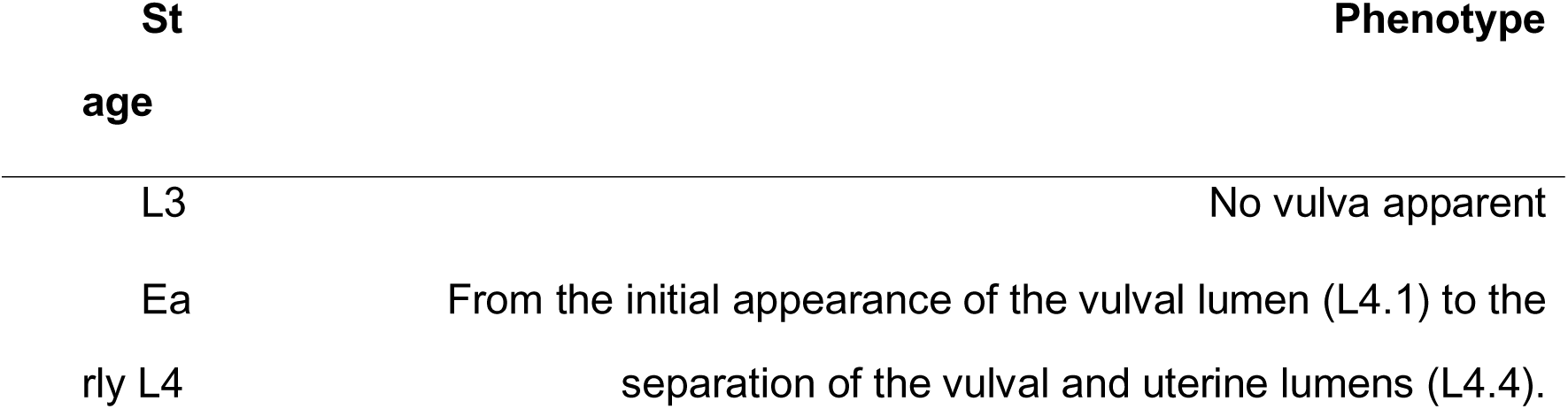
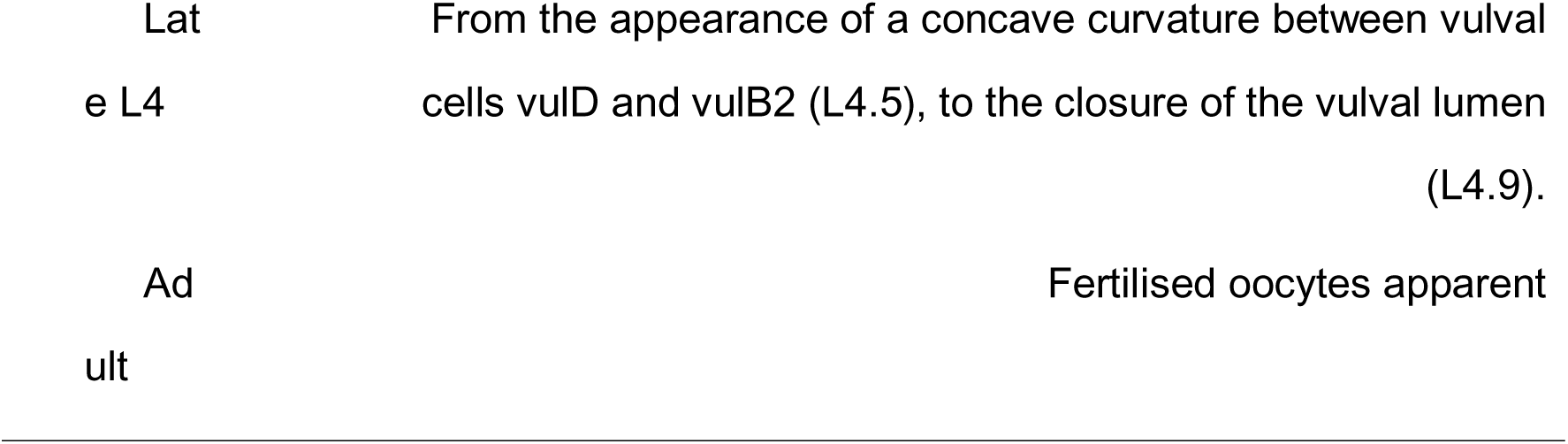
Developmental scoring criteria. Further details can be found in.

### Motility assessment

Manual motility analysis was performed by transferring single worms into 100 µL droplets of M9, waiting for 30 seconds, and counting the number of thrashes (defined as a deviation of the head and tail from the long axis of the body) for next 30 seconds. Automated motility analysis was performed using the MBF Biosciences Worm Lab Imaging System. Nematodes were transferred into 50 µL droplets of M9 placed in the centres of freshly poured NGM plates, and one-minute videos were taken to assess their swimming dynamics. Thrashing rates were quantified by extracting the total number of body bends (> 20° deviation from the long axis of the body) over the length of the footage. Manual and automated thrashing rate measurements were compared using a subset of the total captured footage and were found to be closely correlated (Pearson’s correlation: r = 0.99, P < 0.0001). Alternative metrics of swimming behaviour were calculated directly from the swimming statistics summary provided by the MBF Biosciences Worm Lab imaging software as described^17^.

### Burrowing assay

Pluronic F-127 (Sigma-Aldrich P2443) was prepared at 26% w/w in water and stored at 4°C until fully dissolved. When in use, this pluronic solution was kept on ice to prevent it from solidifying. A 25μl drop of pluronic gel was added to the middle of an empty 3.5cm plate and allowed to set for 5 minutes. 20-25 synchronised worms were then transferred into the drop using an eyelash pick. 6ml of pluronic gel was added on top and allowed to set for 10 minutes, achieving a gel height of ∼0.7cm. A 25ul drop of fresh OP50 culture was added on to the centre of this gel above the initial worm drop. The plate was monitored every 15 minutes for >4h with worms that had burrowed up onto the OP50 drop counted and removed^48^.

### Defecation rates

Defecation rates were assessed as described^49^. Individual nematodes were transferred to fresh NGM plates and allowed to acclimate for five minutes. Following acclimation and the first peristaltic contraction of the body wall muscle, the time between four defecation motor programmes (DMPs) was measured. If no such contraction was observed over a five-minute period, the animal was classified as a non-defecator (0 DMPs min^-1^). Observations were conducted by eye using a Leica S6 E Stereo Zoom light microscope. Only nematodes observed to be in the process of feeding were assayed.

### Pharyngeal pumping

Nematodes were transferred individually to freshly seeded NGM plates and allowed to acclimate for one minute. Following acclimation, the animals were manually observed for 20 seconds using a Leica S6 E Stereo Zoom light microscope and the number of pharyngeal pumps counted.

### Electron microscopy

Nematodes were individually transferred into small quantities of M9 using an eyelash pick, before being fixed in 2.5 % glutaraldehyde fixative in 100 mM sodium cacodylate buffer (CAB; pH 7.2). The heads and tails were separated from the bodies using a scalpel, and bodies were left overnight in fixative at 4 °C. The samples were subsequently washed twice with CAB and re-suspended in 2 % low melting-point agarose in CAB. The worm bodies were identified in the agarose suspension using a Leica S6 E Stereo Zoom light microscope, excised, transferred to glass vials, and stained with 1 % osmium tetroxide in CAB for 1 h at room temperature. Excess stain was removed by washing twice with Milli-Q water, each wash lasting 10 minutes. Subsequently, worms were dehydrated in an ethanol series (50 %, 70 %, 90%, and 100 %) for 10 minutes each, with the 100 % ethanol dehydration step repeated 3 times. Dehydration was followed by two 10-minute washes in propylene oxide, after which the samples were treated with a 1:1 mixture of low-viscosity (LV) resin and propylene oxide for 30 minutes at room temperature. Samples were then transferred to fresh LV resin twice for 2 hours each before being embedded in LV resin via polymerisation at 60 °C for 24 h. Embedded samples were examined under a Leica S6 E Stereo Zoom light microscope to identify and excise the worm bodies, which were then oriented on a resin block for optimal sectioning. 70 nm-thick transverse sections were cut, approximately equidistant from the vulva and the anterior tip of the worm body, using a Diatome diamond knife and an EM UC7 ultramicrotome. Sections were collected onto 400-mesh copper grids and counterstained with 4.5 % uranyl acetate for 45 minutes, followed by Reynolds’ lead citrate for 7 minutes. Sections were imaged using a Jeol 1230 transmission electron microscope with an accelerating voltage of 80 kV, fitted with a Gatan One View 4K digital camera.

### Epifluorescence imaging

Fluorescence imaging of the body wall muscles was performed using GFP-tagged myosin (RW1596 *myo-3 (st386); StEx30 [myo-3p::GFP::myo-3 + rol-6 (su1006)*) and mCherry-tagged actin (WX8490 *YqIs100[ced-1p::mCHERRY::ACT1]*). Nematodes were anaesthetised in 25 mM tetramisole and mounted on 2.5 % agarose pads for imaging using a Leica DMR compound epifluorescence microscope fitted with a 40 X objective. All images of the body wall muscles were captured between the pharynx and vulva.

### Myosin filament density

Myosin filament density was estimated from electron micrographs by manually counting the number of thick filaments in 10 0.65 x 0.52 (length x width) µm cross sections per replicate, with each section spanning the centre of the sarcomere. Cross sectioning was performed blind using the File Name Encrypter tool within the Blind Analysis tool suite for Fiji v. 2.15.1^50^.

### Body wall muscle organisation

Sarcomere disorganisation was assessed qualitatively using a scoring system^16^. Cells received a score of 1 if all filaments were arranged in parallel, symmetric rows; a score of 2 if the filaments were largely arranged in parallel, but contained some gaps; 3 if the filaments appeared crooked or frayed, with an elevated number of gaps between filaments; and 4 if the filaments appeared mostly crooked, broken, and poorly arranged, with substantial gaps between filaments.

### Quantitative measurements of body wall muscle cell area and fibre length

Quantitative measurements were of body wall muscle were performed as previously described^21^. Briefly, muscle:area ratio (gap to total cell area) was calculated from images that included at least on complete muscle cell. Gaps left by degenerating muscle fibres and the total area of a single muscle cell were drawn and calculated in Fiji using the polygon selection tool.

### Muscular contractibility

Muscular contractibility was assessed using the acetylcholine esterase inhibitor levamisole. Groups of 10 nematodes were transferred in technical triplicate to unseeded 30 mm NGM plates and immediately imaged using an MBF Biosciences Worm Lab Imaging System. Immediately after imaging, the animals were transferred to freshly prepared NGM plates containing levamisole (75 µM). Animals were imaged after 10 minutes, and body lengths were quantified in both sets of images using the segmented line tool for Fiji v. 2.15.1. Muscular contractibility was then calculated as the plate-averaged % reduction in body length following levamisole treatment:

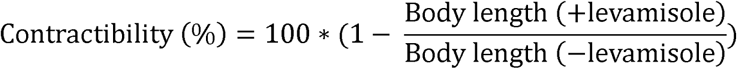

### Mitochondrial morphology and ultrastructure

Mitochondrial morphology was estimated from electron micrographs by manually segmenting all observable mitochondria using the segmented line tool for Fiji v. 2.15.1^50^. To quantify mitochondrial shape, a circularity index (CI) was calculated from area and perimeter measurements as follows:

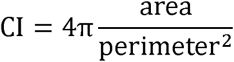

Fluorescence imaging the body wall muscle mitochondrial networks was performed using mitochondrially-targeted GFP under the control of the *myo-3* promoter (SJ4103 *zcls14 myo-3p::GFP[mit]*). Nematodes were anaesthetised in 25 mM tetramisole and mounted on 2.5 % agarose pads for imaging using a Leica DMR compound epifluorescence microscope fitted with a 63 X objective. All images were captured approximately equidistant from the pharynx and vulva.

The integrity of the mitochondrial networks was assessed qualitatively. Each cell received a score based on the integrity of the mitochondrial networks, with cells receiving a score of 1 if the network was entirely tubular and linear; 2 if the network was mostly linear with few spherical mitochondria; 3 if the network appeared mostly fragmented with spherical mitochondria predominant; and 4 if the network was almost entirely fragmented with large regions devoid of observable mitochondria.

### mtDNA copy number

Germ-free *glp-1* animals cultivated at 25°C were used to quantify mtDNA copy number (mtDNA-CN). mtDNA-CN was measured using an adapted one-worm quantitative PCR (qPCR) protocol^29^. Individual worms in lysis buffer (30 mM Tris pH 8, 8 mM EDTA, 100 mM NaCl, 0.7 % NP40, 0.7 % Tween 20, 100 µg mL^-1^ Proteinase K) were centrifuged briefly, and lysed (60 min at 65 °C, 15 min at 95 °C). PCR reactions were assembled in technical duplicate with mtDNA and nDNA (nuclear DNA) primers (Table 2) using a SYBR Green PCR Master Mix (Thermo Fisher), with 2 µL template DNA and a total reaction volume of 25 µL. PCR reactions were cycled in a QuantStudio 3 Real-Time PCR System (2 mins at 50°, 10 mins at 95°C, 40 cycles of 15 sec at 95°C, 60 sec at 62 °C). Relative mtDNA-CNs were calculated using mitochondrial and nuclear-target cycle times (CTs) obtained from the QuantStudio Design and Analysis software, where:

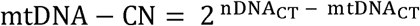

**Table 2.**
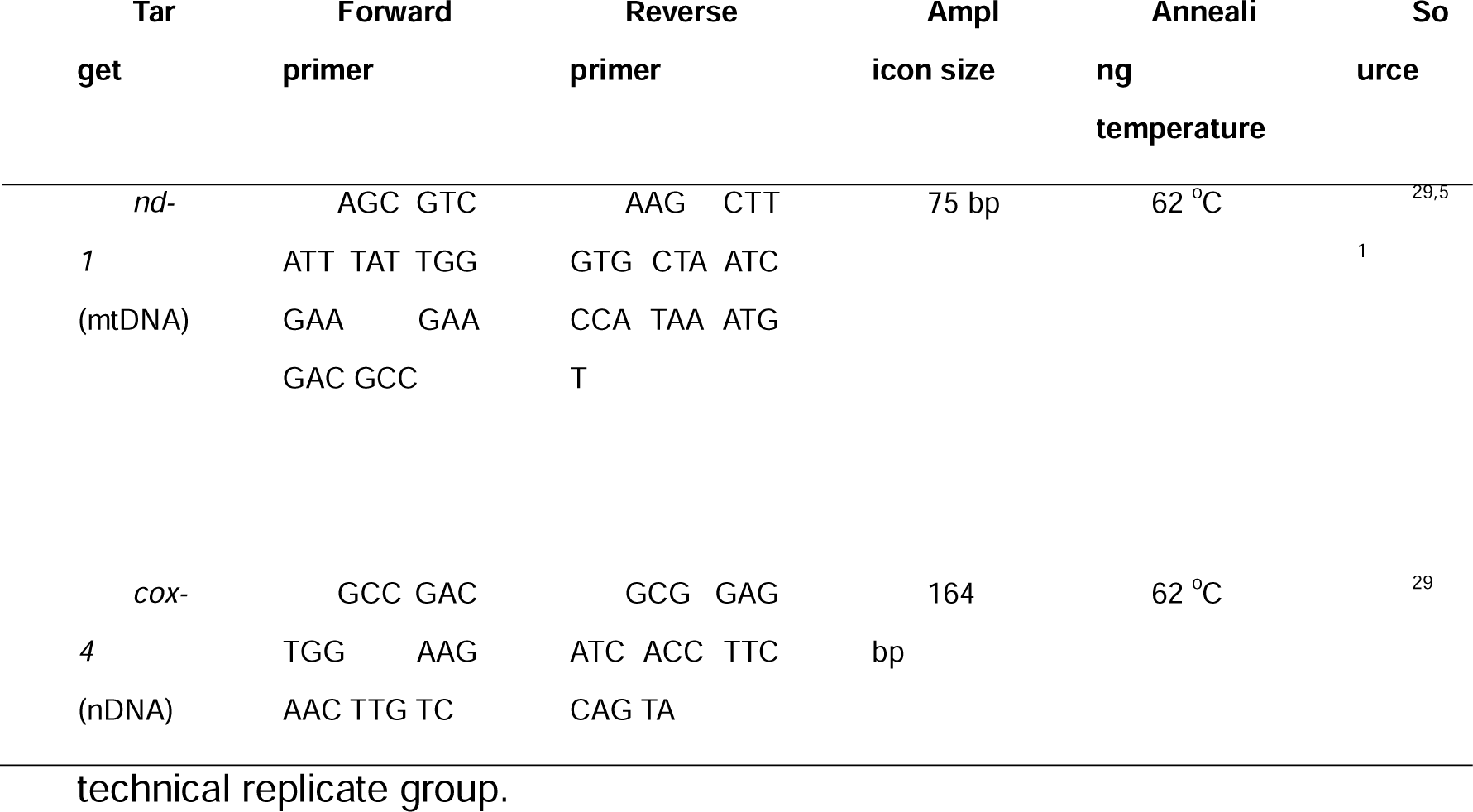
Descriptions of mitochondrial and nuclear primer pairs.

Data was normalised to average relative mtDNA-CNs of relevant *E. coli* OP50-fed controls. Samples were excluded from copy number calculations if the CT standard deviation exceeded 1 for either nDNA or mtDNA primer pair within each

### Whole body ATP measurements

Whole-body ATP levels were assessed as described^30^. Briefly, worms were transferred in groups of five to 50 µL M9 in thin-walled PCR tubes. The samples were then lysed at 95 ^O^C for 15 minutes, incubated at 25 ^O^C for 5 minutes, and assessed using a luminescence-based CellTiterGlo 3D Cell Viability Assay (Promega) in accordance with the manufacturer’s instructions. ATP levels were quantified using a standard curve of ATP standards and were normalised to the average ATP levels of relevant *E. coli* OP50-fed controls. Luminescence measurements were taken using a BMG FLUOStar Omega plate reader.

### ATP measurements in body wall muscle using Queen-2M

For estimates of relative ATP levels using the Queen-2m sensor, λ_ex_ of 400 and λ_em_ of 545 was measured using a Zeiss LSM880 confocal laser scanning microscope with a 40X objective and 400/494 excitation ratio calculated, as described^30,31^. Images were captured as tile-scanned z-stacks through the entire range of observable fluorescence (40-70 µm) and stitched automatically with Zen Blue (Zeiss). The number of slices per stack were determined using the optimal sectioning tool of Zen Blue, with a 50 % overlap between slices. The resulting scans were then maximally projected for image analysis, and relative ATP levels were calculated by dividing the mean whole-body fluorescence at 405 nm λ_ex_ by the mean whole-body fluorescence at 488 nm λ_ex_. These ratios were then converted to relative ATP levels by normalising to the average 405ex/488ex ratios of relevant *E. coli* OP50-fed controls.

### Statistics

Statistical analysis was performed using GraphPad Prism 10 and R v. 4.3.3. All age-related data were analysed by two-way ANOVA, with age and bacterial diet included as an interaction term. Post-hoc comparisons were performed using Student’s t tests. Corrections for multiple comparisons were conducted where appropriate using the False Discovery Rate (FDR) method. F tests were used to compare variances. Unless stated otherwise, all remaining data were analysed using Student’s t tests.

---

**Figure S1.**
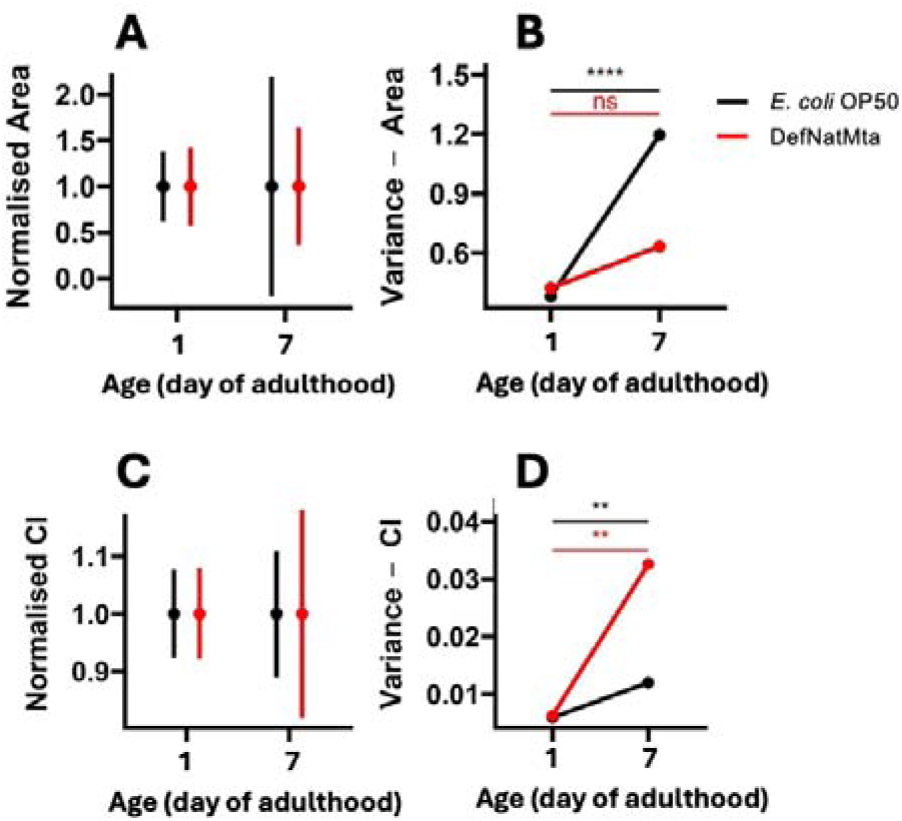
**A-B)** Normalised average mitochondrial area ± variance (A) and mitochondrial area variance (B) calculated from transverse sections represented in Fig. 3C-D. **C-D)** Normalised average mitochondrial CI ± variance (C and mitochondrial CI variance (D) calculated from transverse sections represented in Fig. 3C-D. Mitochondrial ultrastructure and morphology was measured in all observable mitochondria per transverse section: n = 23-129 pooled from 2 experiments. Significances presented represent variance comparisons between day 1 and day 7 using Fisher’s F-tests. ***, P < 0.001; **, P < 0.01; ns, not significant (P > 0.05).

**Figure S2.**
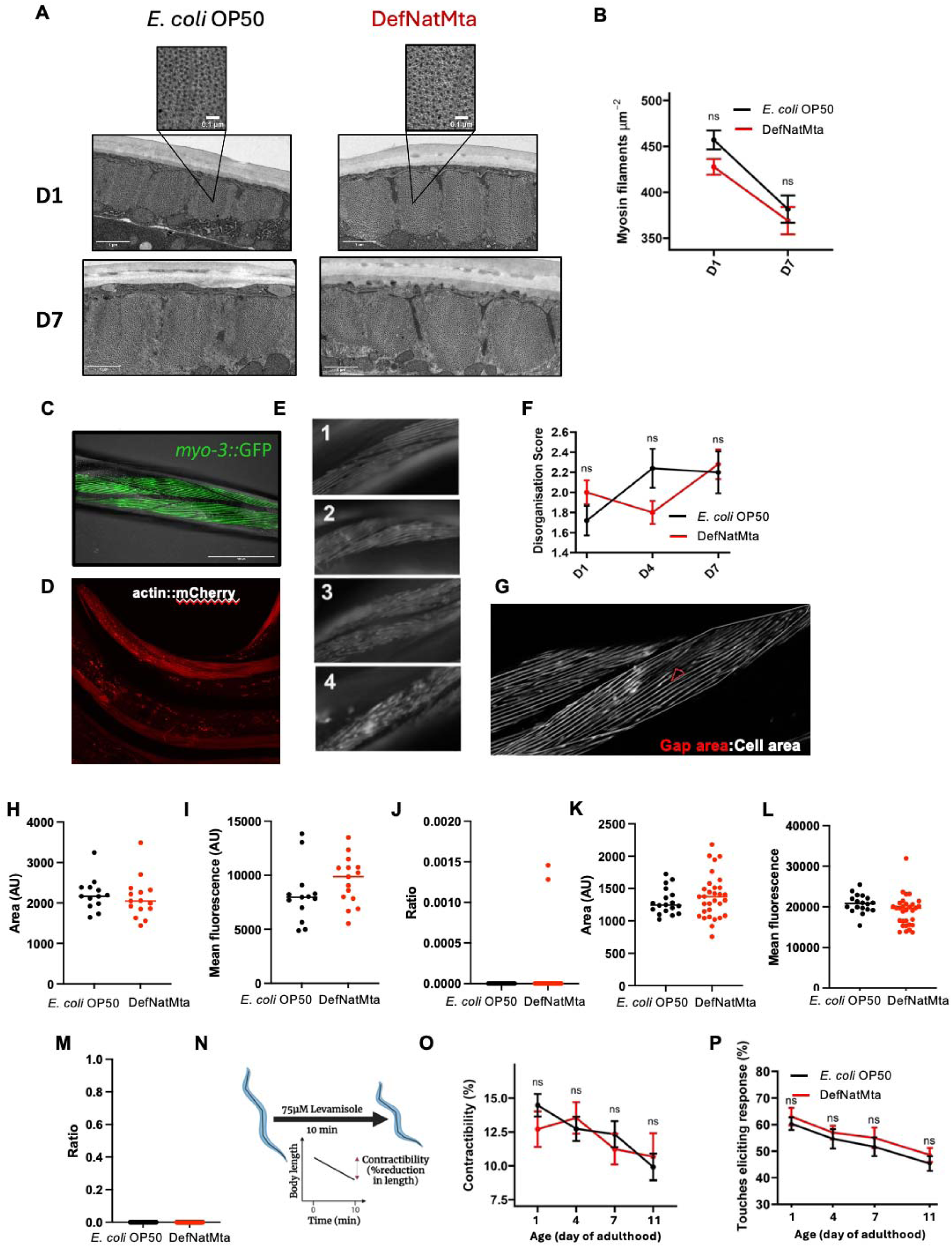
DefNatMta does not impact on host body wall musculature and neuromuscular transmission. **A**) Representative electron micrographs of body-wall muscle blocks from OP50- and DefNatMta-fed worms. Insets are representative cross-sections used for assessing myosin filament density. Scale bars (main images): 1 µm; Scale bars (insets): 100nm. **B**) Myosin filament densities of OP50- and EM-fed worms, assessed from cross-sections (as in F, insets) of randomly chosen muscle blocks. N = 20 pooled from 2 experiments. **C**) Representative image of muscle cells expressing *myo-3::GFP* (RW1596 *myo-3 (st386); StEx30[myo-3p::GFP::myo-3 + rol-6 (su1006)]*), showing the anterior half of the worm. **D**) Representative image of muscle cells expressing mCherry-tagged actin. **E**) Representative scoring system for assessing sarcomere disorganisation. 1 = Fibres organised in parallel, symmetric rows; 2 = Fibres mainly parallel, containing some gaps; 3= Fibres lie in same direction but contain gaps, bends, are frayed; 4 = Fibres are broken and bent with a ‘blurred’ appearance. **F**) Sarcomere disorganisation scores in OP50 and EM-fed worms. N = 25-27 (1 experiment). **G**) Example image of how the muscle area ratio (gap to total cell area) was calculated. Gaps left by degenerating muscle fibres (red-lined area) and the total area of a single muscle cell (white-lined area) were drawn and calculated in Fiji using the polygon selection tool. **H-J**) Measures GFP-tagged myosin in single muscle cells. Area covered (H), mean fluorescence (I), Gap:cell area ratio (J). N = 13 **K-M**) Measures mCherry-tagged actin in single muscle cells. Area covered (H), mean fluorescence (L), Gap:cell area ratio (M). N = 35 pooled from 2 experiments. **N**) Schematic of levamisole treatment contractibility assay. Images of worms were taken before and after levamisole treatment, and contractibility was defined as the % reduction in average worm length following treatment. **O**) Contractibility of OP50- and DefNatMta-fed worms following levamisole treatment. N = 60 pooled from 2 experiments with three technical replicates. **P**) Touch-responsiveness of OP50- and EM-fed worms. N = 21-30 pooled from 3 experiments.

## Notes

### Competing Interest Statement

The authors have declared no competing interest.

